# Functional characterization of odor responses and gene expression changes in olfactory co-receptor mutants in *Drosophila*

**DOI:** 10.1101/2021.06.18.449017

**Authors:** Alina Vulpe, Pratyajit Mohapatra, Karen Menuz

## Abstract

Two large families of olfactory receptors, the Odorant Receptors (ORs) and the Ionotropic Receptors (IRs), mediate responses to most odors in the insect olfactory system. Individual odor binding “tuning” OR receptors are expressed by olfactory neurons in basiconic and trichoid sensilla and require the co-receptor Orco to function. The situation for IRs is more complex. Different tuning IR receptors are expressed by olfactory neurons in coeloconic sensilla and rely on either the Ir25a or Ir8a co-receptors; some evidence suggests that Ir76b may also act as a coreceptor, but its function has not been systematically examined. This is particularly important as recent data indicate that nearly all coeloconic olfactory neurons co-express Ir25a, Ir8a, and Ir76b. Here, we report the effects of *Drosophila* olfactory co-receptor mutants on odor detection by coeloconic olfactory neurons and determine their broader impact on gene expression through RNASeq analysis. We demonstrate that Ir76b and Ir25a function together in all amine-sensing olfactory receptor neurons. In most neurons, loss of either co-receptor abolishes amine responses, whereas in ac1 sensilla, amine responses persist in the absence of Ir76b or Ir25a, but are lost in a double-mutant. Such responses do not require Ir8a. Conversely, acid-sensing ORNs require Ir8a, but not Ir76b or Ir25a. Using antennal transcriptional profiling, we find that the expression of acid-sensing IR receptors is significantly reduced in *Ir8a* mutants, but is unaffected by the loss of *Ir25a* or *Ir76b.* Similarly, select OR tuning receptors are also downregulated in *Orco^2^* mutants. In contrast, expression of amine-sensing IR receptors is mostly unchanged in *Ir25a* and *Ir76b* mutants. Together, our data reveal new aspects of co-receptor function in the olfactory system.

**Summary:** Insect vectors of human disease rely on their sense of smell to locate humans for blood meals. A critical first step in olfaction is the odorant-induced activation of receptors on olfactory neurons. There are two major olfactory receptor families in insects, with each species having dozens of different odorant-binding “tuning” receptors. The receptor complexes also contain non-tuning co-receptors, which are highly conserved across insect species and are required for function. Here we characterize co-receptor mutants with electrophysiological recordings and transcriptome analysis in *Drosophila*. Our findings resolve the differential co-receptor dependence of olfactory neuron responses to volatile amines and acids. We also report changes in antennal gene expression that result from the absence of these co-receptors. Most notably, the absence of some co-receptors leads to a selective loss of transcript expression for the tuning olfactory receptors whose function depends on the missing co-receptors. Together our data provide new insight into the roles of co-receptors in insect olfaction.

## Introduction

A vast number of different volatile molecules can be detected as odorants by both vertebrate and invertebrate animals. In insects, nearly all odorants are detected by receptors in one of two large gene families, the Odorant Receptors (ORs) and the Ionotropic Receptors (IRs) [1, 2]. These receptors are found on the dendrites of olfactory receptor neurons (ORNs) housed in antennal olfactory sensilla. Large scale screens in *Drosophila* have shown that IR receptors generally detect polar compounds such as acids and amines, and they are expressed by ORNs in coeloconic sensilla [3, 4]. OR odorant receptors respond to alcohols, ketones, and aldehydes as well as pheromones; nearly all are expressed by ORNs housed in basiconic or trichoid sensilla [5–10]. Most ORNs express a single type of “tuning” Or or IR that determines the odorants to which the receptors can bind.

ORs and IRs form ligand-gated non-selective cation channels, which consist of tetrameric complexes [11–13]. The Odorant Receptor Co-Receptor, Orco, pairs with the different OR tuning receptors to form functional receptors and is required for the trafficking of OR tuning subunits to olfactory neuron dendrites [11, 14]. Accordingly, loss of Orco function leads to the elimination of odor responses in OR-expressing ORNs [14]. In *Drosophila*, age-related ORN degeneration occurs in the absence of Orco, although it is unclear whether this occurs to all or subsets of Orco^+^ neurons [15–17]. Orco is highly conserved, unlike most tuning ORs, and mutation of Orco similarly impairs olfactory signaling and neuronal health in other insect species [18–23].

There are three potential co-receptors for IR receptors: Ir25a, Ir8a, and Ir76b. In *Drosophila*, each of the four functional classes of coeloconic sensilla contain one ORN that detects acidic odors, and these responses require the co-receptor Ir8a [3, 12, 24]. An acid sensing ORN population in the sacculus also relies on this co receptor [25]. Similar to Orco, the function of Ir8a is conserved in other insect species [26, 27]. However, it is unknown whether neuronal loss also occurs in the absence of Ir8a or other IR co-receptors.

Other ORNs in coeloconic sensilla respond to amines [3, 24], but here the co-receptor requirements are less clear. It was first reported that Ir25a is the required co-receptor for amine-sensing IRs based on the loss of amine-responses mediated by ORNs in ac2 and ac4 sensilla in *Ir25a* mutants [12]. However, later studies found that amine responses mediated by ac2 Ir41a^+^ ORNs require Ir76b, but not Ir25a, and that no IR co-receptors are required for amine responses mediated by ac1 Ir92a^+^ ORNs [28, 29]. Reconstitution experiments using ac4 amine-receptor Ir76a demonstrated that both Ir25a and Ir76b are needed to form functional receptors [12], similar to the joint requirement for Ir25a and Ir76b in IR receptors that detect fatty acids, sour and carbonation in the gustatory system [30–32]. However, coeloconic ORN amine sensitivity has not been systematically examined in Ir76b mutants, and the co-receptors for amine sensing neurons are still unresolved.

Our understanding of co-receptor function has been further challenged with the report of new co-receptor knock-in reporter lines that reveal much broader expression of co-receptors than originally anticipated [33, 34]. In *Drosophila*, most coeloconic ORNs express Ir8a, Ir76b, and Ir25a [34]. Ir25a was additionally found in most OR-expressing basiconic and trichoid neurons, although no tuning IRs are known to be expressed in these neurons.

Here we examined the function of IR co-receptors in *Drosophila* olfaction using single-sensillum electrophysiology and transcriptional profiling of co-receptor mutants. We discover that amine responses in all coeloconic sensilla rely on Ir25a and Ir76b, including responses mediated by a previously undetected amine-sensing IR in ac3B ORNs. Further, expression of acid-tuning IRs is greatly reduced in the absence of Ir8a, as is the expression of many Ors in an Orco mutant.

## Results

### Both Ir25a and Ir76b are co-receptors for amine sensing neurons

We first examined the co-receptor dependence of responses to amines in ac2 and ac4 sensilla, which are mediated by ORNs expressing Ir41a (ac2), Ir76a (ac4), and Ir75d (ac2 and ac4) [3]. We performed single-sensillum recordings (SSR) to compare odor-induced spiking responses in sensilla from co-receptor mutants with those from WT flies. We quantified the summed activity of all neurons in each sensilla due to the difficulty of reliably sorting spikes in coeloconic neurons [3]. For this reason, we selected odorants known to primarily activate specific neurons. We found that responses to both pyrrolidine (Ir75d) and 1,4-diaminobutane (Ir41a) were eliminated in both Ir25a and Ir76b mutants in ac2 sensilla (Figure 1A). Likewise, excitatory responses in ac4 sensilla to phenethylamine (Ir76a), pyrrolidine (Ir75d), and ammonia (Ir76a) were lost in both Ir25a and Ir76b mutants (Figure 1B). Responses were not reduced in Ir8a mutants, although they were somewhat larger than the control line for some sensilla and odor combinations. Together, this suggests that only Ir25a and Ir76b are necessary for proper function of these amine receptors, despite co-expression of Ir8a in these neurons.

**Figure 1.**
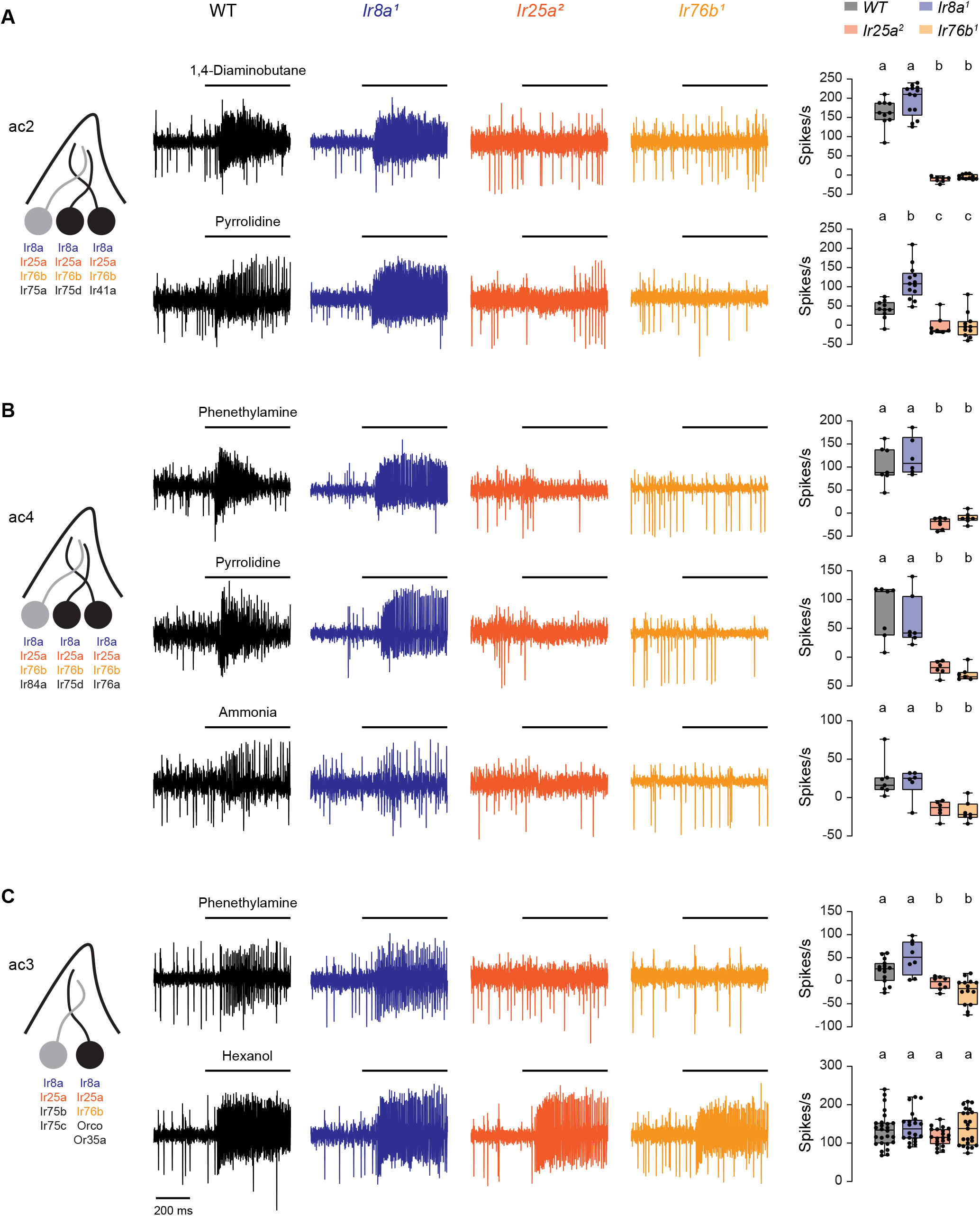
Both Ir25a and Ir76b are required co-receptors for amine-sensing receptors in ac2-4 sensilla. (A) Left, diagram of ac2 sensilla, showing receptor expression in each ORN. ORNs that respond to the tested amines are in black. Middle, representative traces of responses to 1,4-diaminobutane and pyrrolidine in ac2 sensilla in WT, *Ir8a^1^*, *Ir25a^2^*, and *Ir76b^1^* fly lines. Right, box plots overlaid with individual data points (n=7-13 sensilla). Genotypes that are significantly different are indicated with different letters. (B) Similar to (A), but for ac4 sensilla and phenethylamine and pyrrolidine (n=6-7 sensilla). (C) Similar to (A), but for ac3 sensilla and phenethylamine and hexanol (n=8-26 sensilla).

In addition to ac2 and ac4, we further examined ac3 sensilla. These sensilla house two ORNs: ac3A, an acid sensitive neuron, and ac3B, a neuron that expresses Orco and a tuning receptor Or35a that together mediate detection of nearly all odors by this neuron [3, 24]. The ac3B ORN additionally expresses Ir8a, Ir25a, and Ir76b, but their function is unclear as this ORN is not known to express a tuning receptor from the IR family [3, 4, 34]. We found that responses to phenethylamine were mediated by ac3B based on its characteristically small spike amplitude and were reduced in *Ir25a* and *Ir76b* mutants (Figure 1C). This is consistent with the persistence of phenethylamine sensitivity in *Orco* mutants [3], and suggests that an unidentified IR tuning receptor is expressed in ac3B ORNs. If so, signaling by OR and IR receptors within ac3B ORNs is functionally independent because responses to hexanol, an odor detected by Or35a/Orco receptors, are unaffected by the loss of any IR co-receptors (Figure 1C).

### Either Ir25a or Ir76b is sufficient to maintain amine responses mediated by Ir92a

We also examined the co-receptor dependence of amine responses in ac1 sensilla. These sensilla contain four neurons, three of which respond to amines or to the related compound ammonia (Figure 2A). One neuron expresses Ir75d, similar to ac2 and ac4 sensilla, whereas Ir92a ORNs respond to a large variety of amines [3, 29, 35]. Intriguingly, Ir92a signaling was shown to be independent of Ir25a, Ir76b, and Ir8a, despite the expression of all three co-receptors in these neurons [4, 29, 34]. We verified this co-receptor independence when measuring ac1 responses to trimethylamine and 1,4-diaminobutane, two amines that activate Ir92a ORNs (Figure 2B) [29, 35].

**Figure 2.**
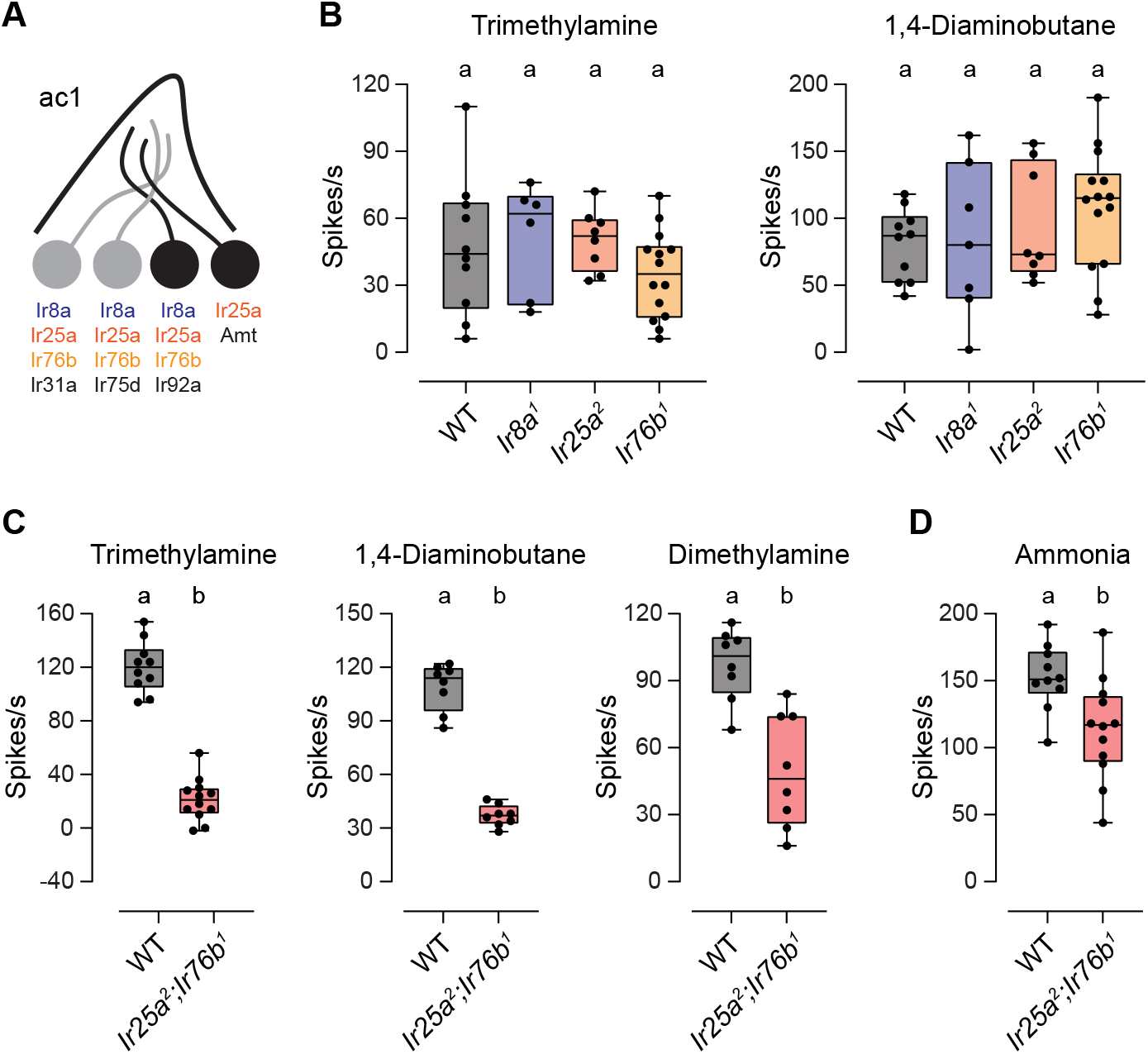
Ir25a and Ir76b play a redundant role supporting the function of the ac1 amine receptor. (A) Diagram of ac1 sensilla, showing receptor expression in each ORN. Those neurons that respond to the tested odors are in black. (B) Box plots showing the distribution of responses to trimethylamine and 1,4-butanedione in ac1 sensilla in WT, *Ir8a^1^*, *Ir25a^2^*, and *Ir76b^1^* flies (n=6-14 sensilla). (C) Box plots showing the distribution of responses to trimethylamine, 1,4-butanedione, and dimethylamine in ac1 sensilla in WT and *Ir25a^2^*;*Ir76b^1^* flies (n=8-12 sensilla). (D) Box plot showing the distribution of responses to ammonia in WT and *Ir25a^2^*;*Ir76b^1^* flies (right) (n=10-12 sensilla). Genotypes that are significantly different are indicated with different letters.

We wondered whether Ir25a and Ir76b play a redundant role in Ir92a neurons. To test this possibility, we generated a double-mutant Ir25a^2^; Ir76b^1^ fly line. Responses to trimethylamine and 1,4-diaminobutane were reduced ~82% and ~66%, respectively, in the absence of these two co-receptors (Figure 2C). Likewise, responses to dimethylamine, a third odor that activates Ir92a ORNs [3, 29, 35], was reduced ~49% in the absence of both Ir25a and Ir76b (Figure 2C). Together our data indicate that all amine-sensing IR receptors including Ir92a rely on Ir25a and Ir76b, with most requiring both co-receptors, and Ir92a able to function with either co-receptor alone.

Two neurons in ac1 sensilla respond to ammonia, those that express Ir92a and those that express Amt, an ammonium transporter that serves as an olfactory receptor [35]. Unlike ac1 responses to amines, ammonia responses were only slightly reduced (~23%) in the absence of both Ir25a and Ir76b (Figure 2D). The reduced portion likely represents the response mediated by Ir92a ORNs because Amt ORNs function independently of IR receptors [35].

### Ir8a is the only co-receptor required for responses in acid sensing neurons

Responses to acidic odors are mediated by olfactory neurons expressing Ir31a (ac1), Ir75a (ac2), Ir75b and Ir75c (ac3), and Ir84a (ac4) [3, 36]. The function of these receptors was reported to require Ir8a, but not Ir25a [12]. However, similar to amine-sensing neurons, recent data indicate that both Ir25a and Ir76b are co-expressed in most of these ORNs (Figure 3) [34]. Given that the role of Ir76b in these neurons has not been tested previously, we decided to examine the Ir76b-dependence of responses in these neurons. We also re-examined the roles of Ir25a and Ir8a.

**Figure 3.**
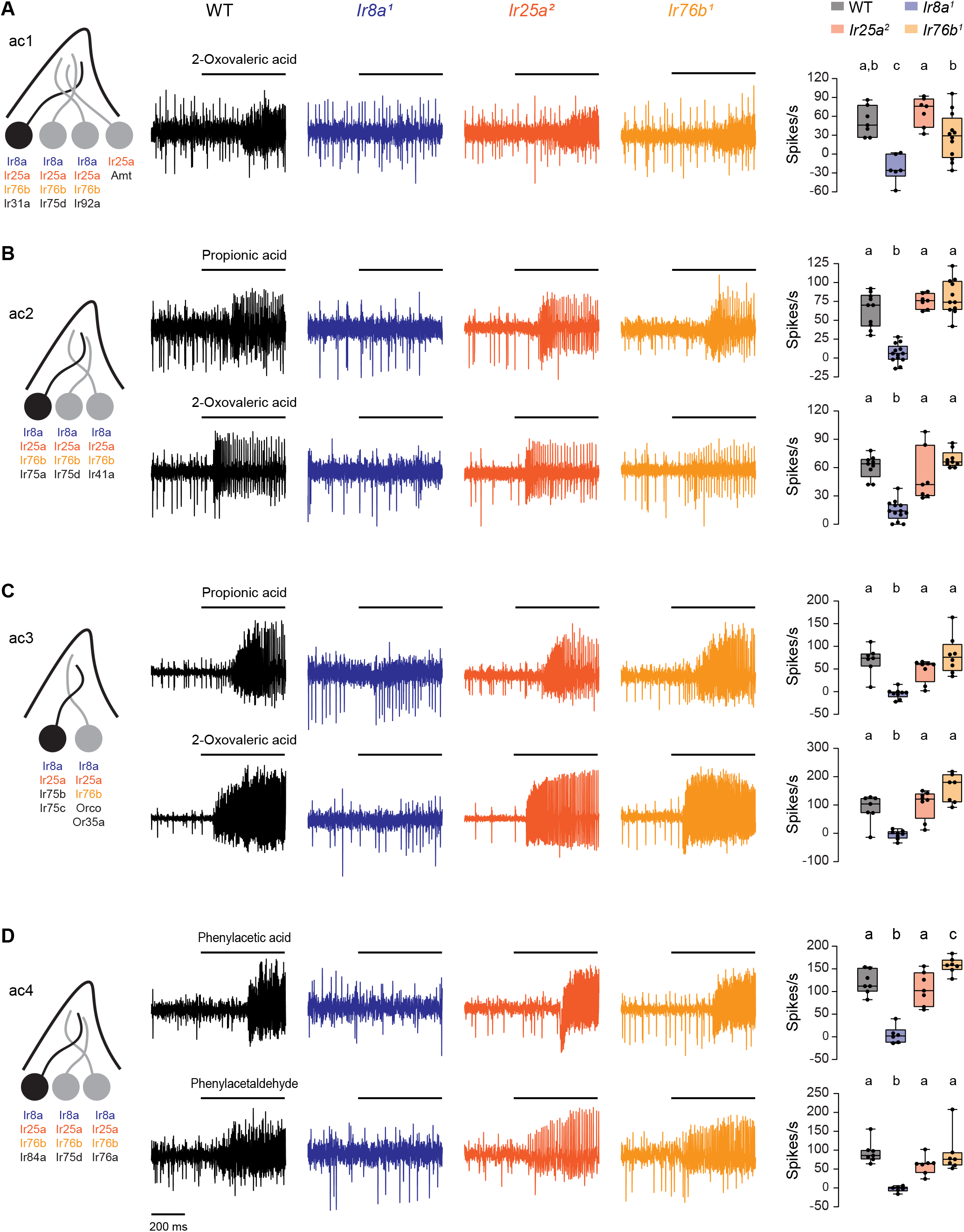
Ir8a is the sole co-receptor required for the function of acid-sensing receptors. (A) Left, diagram of ac1 sensilla and receptor expression. The acid sensing ORN is in black. Middle, representative traces of responses to 2-oxovaleric acid in ac1 sensilla in WT, *Ir8a^1^*, *Ir25a^2^*, and *Ir76b^1^* fly lines. Right, box plots overlaid with individual data points (n=6-12 sensilla). (B) Similar to (A), but for ac2 sensilla and propionic acid and 2-oxovaleric acid (n=7-13 sensilla). (C) Similar to (A), but for ac3 sensilla and propionic acid and 2-oxovaleric acid (n=7-9 sensilla). (D) Similar to (A), but for ac4 sensilla and phenylacetic acid and phenylacetaldehyde (n=6-7 sensilla). Genotypes that are significantly different are indicated with different letters.

We measured SSR responses to odors known to be detected by Ir31a, Ir75a, Ir75b/c, and Ir84a ORNs in co-receptor mutants [3, 37]. Responses to acids in each of the four sensilla were entirely eliminated in *Ir8a^1^* mutant flies, consistent with previous findings (Figure 3). In contrast, responses did not require either IR25a or Ir76b (Figure 3). Thus, Ir8a acts as the sole co-receptor for acid-sensing receptors.

### Transcriptional changes induced by loss of co-receptors

Antennal Or22a^+^ and Or47b^+^ ORNs olfactory receptor neurons undergo axonal and/or somatic degeneration with age in *Orco* mutants, perhaps as a consequence of reduced neuronal activity [15, 17]. We turned to transcriptomics to determine if similar changes occur when IR co-receptor expression is absent, as evaluated by a decrease in tuning IR expression. We also examined whether loss of co-receptors induces broader changes in antennal gene expression. We used two independent samples of antennal RNA from WT, *Ir8a^1^*, *Ir25a^2^*, and *Ir76b^1^* loss-of-function mutants for RNASeq transcriptional profiling (Figure 4A). To compare the effects of IR coreceptor loss with effects induced by loss of the co-receptor Orco from basiconic and trichoid sensilla, we also analyzed samples from *Orco^2^* mutant flies. As expected, the expression level of most genes in each mutant line was similar to WT flies (Figure 4B-E). Expression of Ir8a was entirely eliminated in *Ir8a^1^* mutants (Figure 4B), consistent with the removal of the entire gene region in this line [12]. Similarly, expression of Ir25a and Orco were reduced to <1% of WT levels in *Ir25a^2^* and *Orco^2^* flies, respectively (Figure 4C and 4E). Surprisingly, *Ir76b* expression was only moderately reduced (~44%) in the *Ir76b^1^* mutants (Figure 4D). Closer examination of *Ir76b* expression on an exon by exon basis revealed that expression of the first 3 of 5 exons of the *Ir76b* genes was nearly absent in *Ir76b^1^* mutants (Figure 4F), in agreement with the selective ablation of these exons in this mutant [38].

**Figure 4.**
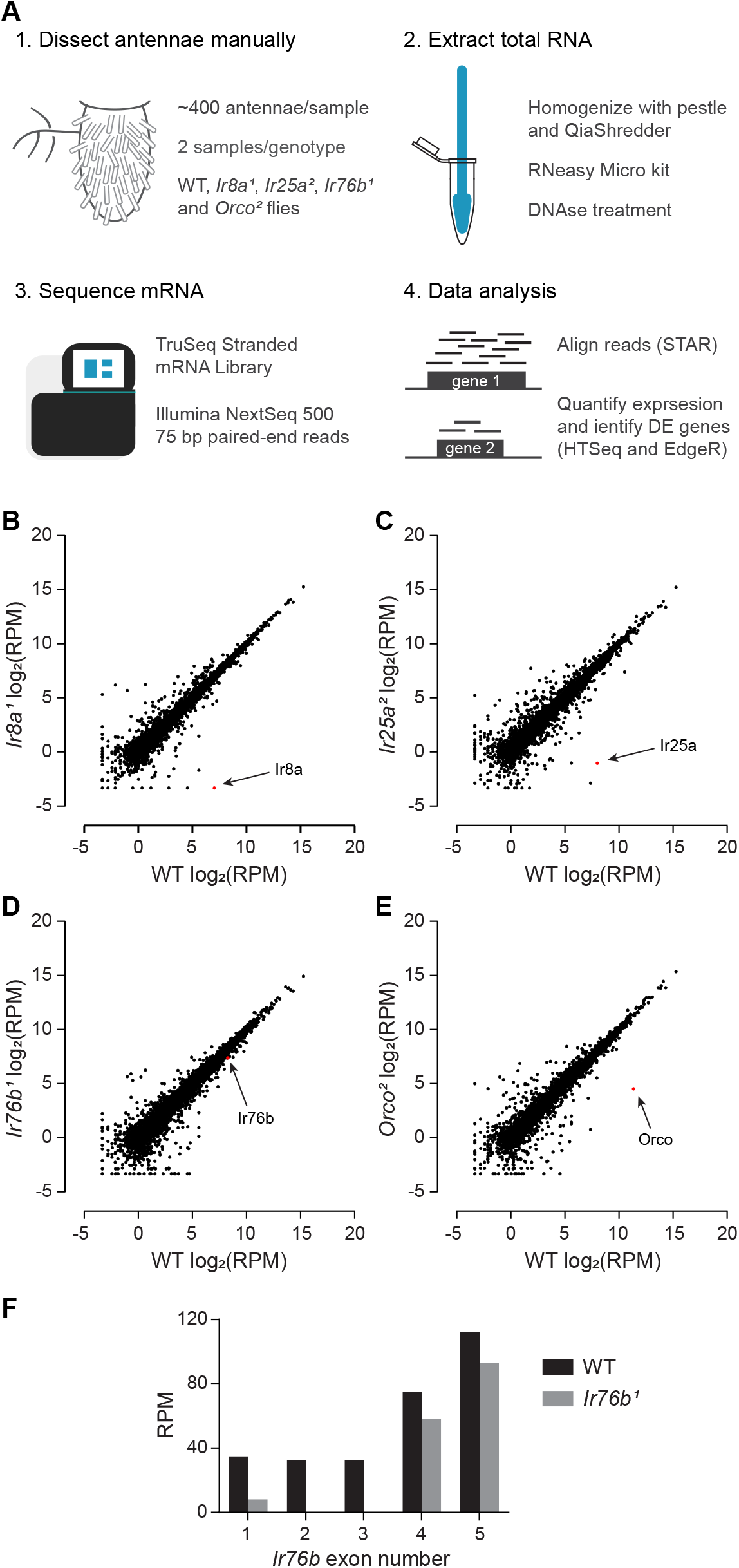
Transcriptional profiling reveals that expression of most genes is unchanged in coreceptor mutants. (A) Workflow for the RNASeq analysis. (B) Expression of each detected gene (black dots) is plotted based on its log_2_ expression in the WT sample and its log_2_ expression in the *Ir8a^1^* mutant. The dot for Ir8a is colored red and labeled with an arrow. (C-E) Similar to (B), but for (C) *Ir25a^2^*, (D) *Ir76b^1^*, (E) and *Orco^2^*. (F) The expression level in RPM for each exon of the *Ir76b* gene in WT flies and *Ir76b^1^* mutants.

We used EdgeR to identify genes that were differentially expressed (DE) between each co-receptor mutant and the WT control (see Methods). There were 233 genes that were differentially expressed in at least one genotype with a false discovery rate (FDR) < 0.05 (Figure 5A-B and Dataset S1). The number of DE genes was highest in *Ir25a^2^* flies (119 genes) and lowest in *Ir8a^1^* (35 genes) (Figure 5C-D). In most genotypes, there are slightly more DE genes that are downregulated than upregulated (Figure 5D).

**Figure 5.**
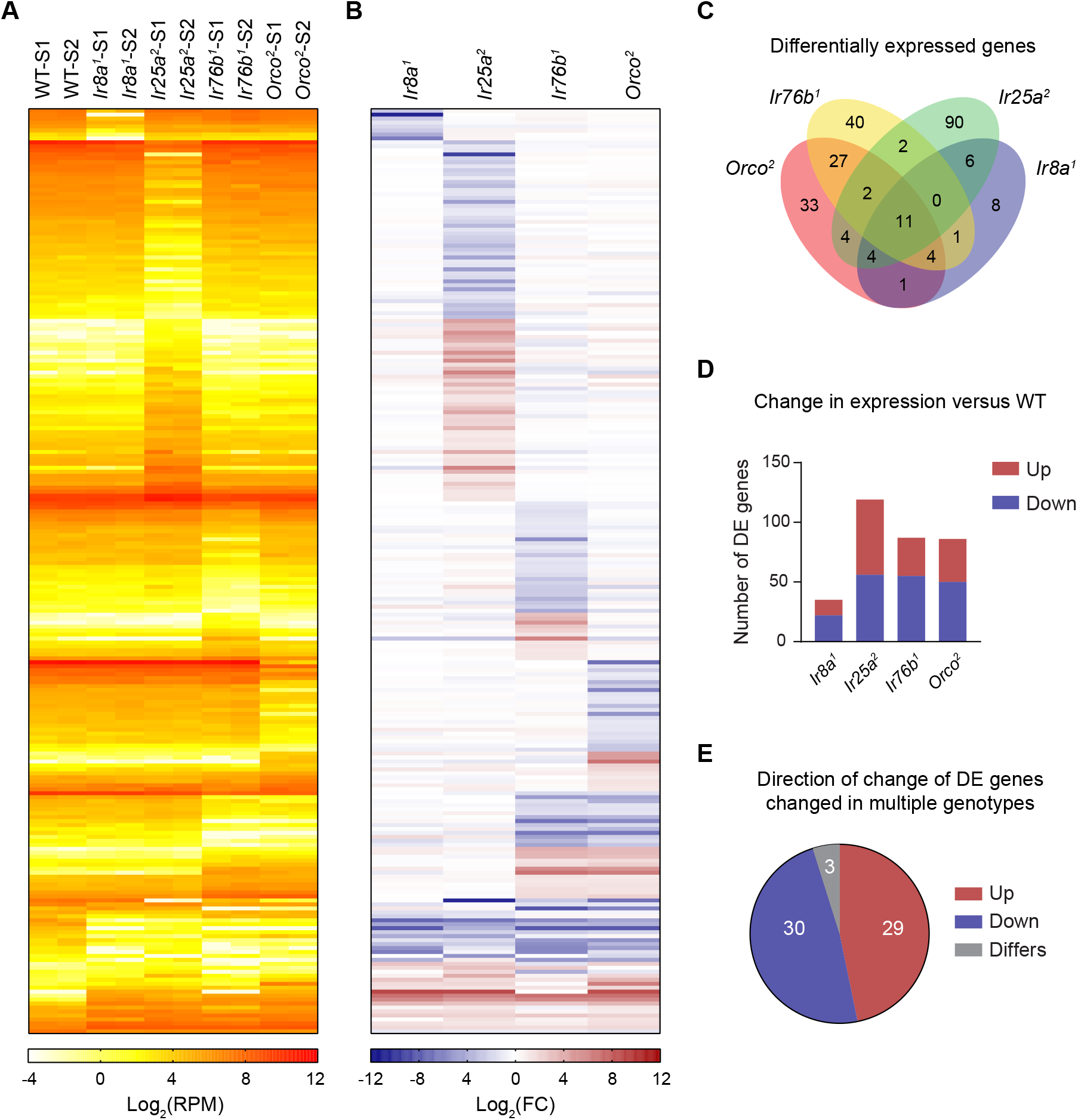
233 genes are differentially expressed in co-receptor mutants. (A) A heat map showing the expression in Log_2_(RPM) of each of the 233 DE genes in each of the ten samples. (B) A heat map reporting the log_2_ of the expression ratio between the co-receptor mutant and WT flies. (C) A Venn diagram showing the number of differentially expressed genes that was changed in each co-receptor mutant or in multiple co-receptor mutants. (D) A bar graph showing the total number of DE genes identified in each co-receptor mutant. The DE genes are shown in maroon if they were detected at higher levels in the mutant than in WT flies, and in blue if they were found at lower levels in the mutant. (E) The pie chart shows the agreement between the direction of change in expression for those DE genes that were DE in multiple genotypes.

The majority of DE genes are changed in a single genotype (171/233, 73%) (Figure 5C and Dataset S1), suggesting that loss of co-receptors predominantly leads to differing transcriptional pathways. We more closely examined the 62 DE genes changed in multiple genotypes. The direction of change (upregulated versus downregulated), was consistent between genotypes (Figure 5E), with only three genes showing differential directions of regulation.

### Selective loss of olfactory receptor expression in co-receptor mutants

We next sought to determine the types of genes whose expression was altered in each coreceptor mutant. PANTHER was used to identify statistically enriched Gene Ontology (GO) terms by quantitatively comparing the proportion of DE genes assigned to particular GO molecular function term versus the proportion among all 13,811 FlyBase genes assigned to at least one GO term (Figure 6). We analyzed the upregulated and downregulated DE genes for each co-receptor mutant separately. No GO terms were enriched among the DE genes upregulated in any of the co-receptor mutants, but several terms were enriched among the DE genes with reduced expression in the absence of particular co-receptors.

**Figure 6:**
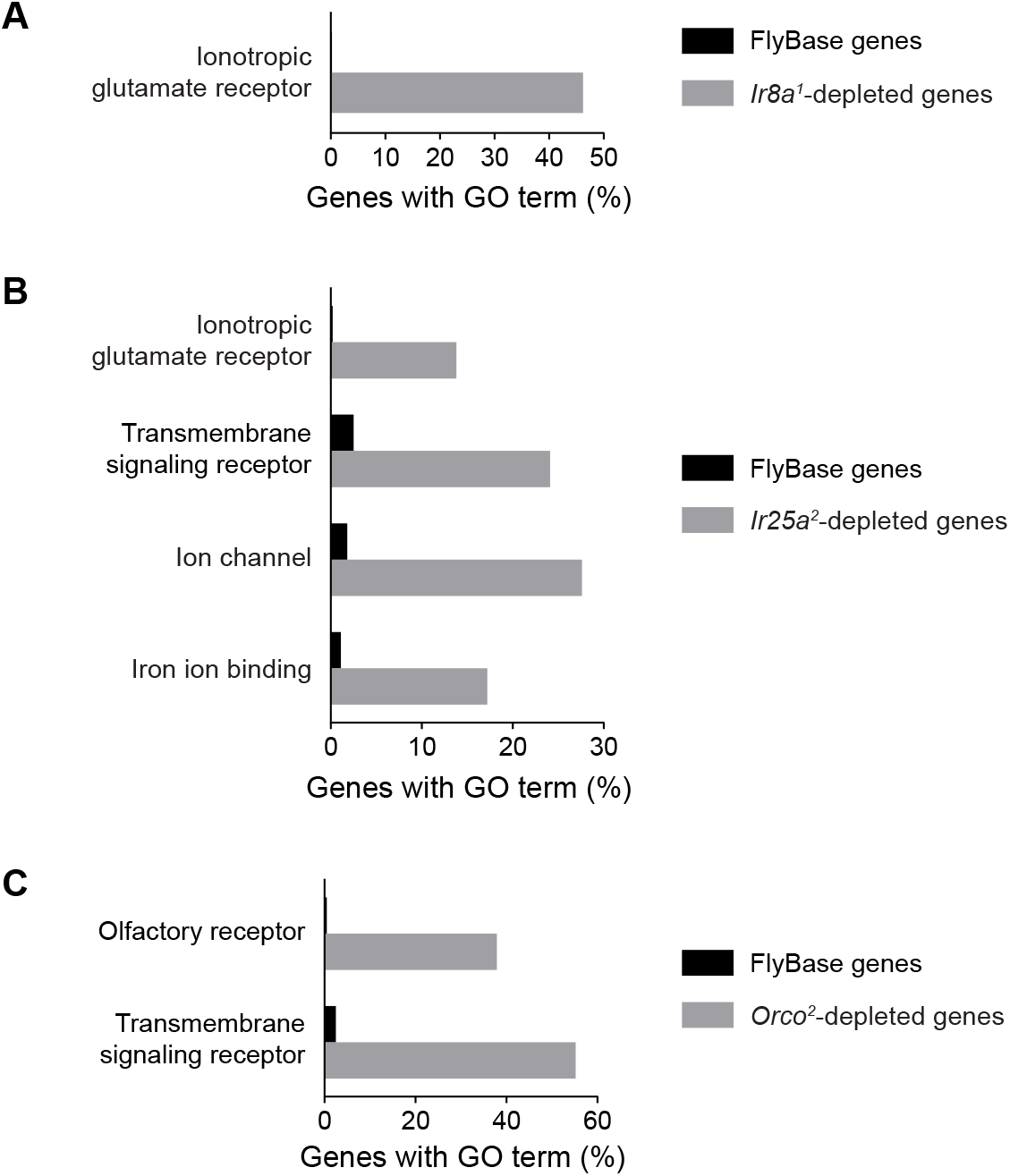
GO analysis identifies molecular function terms enriched amongst genes depleted in co-receptor KOs. (A) The percent of DE genes depleted in *Ir8a^1^* flies with the Gene Ontology (GO) term “ionotropic glutamate receptor” as compared to all Drosophila melanogaster genes listed in FlyBase. (B) The percent of DE genes depleted in *Ir25a^2^* flies with the GO terms “ionotropic glutamate receptor”, “transmembrane signaling receptor”, “ion channel”, and “iron ion binding” as compared to all Drosophila melanogaster genes listed in FlyBase. (C) Similar to (B), but for the DE genes depleted in *Orco^2^* mutants and the GO terms “olfactory receptor” and “transmembrane signaling receptor”.

The most striking finding was the reduced expression of OR and IR receptors in several co-receptor mutants. Genes associated with GO:0004970 “ionotropic glutamate receptor” were statistically enriched among the DE genes with reduced expression in either *Ir8a^1^* flies or *Ir25a^2^* flies (6 and 4 IRs, respectively) (Figure 6A-B). Similarly, GO:0004984 “olfactory receptor” was enriched amongst the DE genes with reduced expression in *Orco^2^* flies (Figure 6C); each of the 11 genes in this category is a member of the OR family. Two additional GO terms, GO:0004888 “transmembrane signaling receptor” and GO:0005216 “ion channel”, were enriched among genes downregulated in *Ir25a^2^* flies (7 and 8 genes, respectively), with the ionotropic receptors forming a large subset of each category. The term “transmembrane signaling receptor” was also enriched among downregulated DE genes in *Orco^2^* flies (16 genes) and consisted primarily of OR olfactory receptors. In contrast to the other co-receptor mutants, no GO terms were enriched among the genes downregulated in *Ir76b^1^* flies.

To gain a more global view of the transcriptional changes in olfactory receptor expression, we inspected the expression levels of all OR and IR receptors known to be expressed in the antenna in each of the co-receptor mutants. Interestingly, nearly all acid-sensing IRs (*Ir75a*, *Ir75b*, *Ir75c*, *Ir64a*, and *Ir31a*) are among the *Ir8a^1^* DE genes, each with <30% of control expression; their expression was indifferent to the loss of other co-receptors (Figure 7A-B). The only other acid-sensing IR, *Ir84a*, had reduced expression with about 56% of the WT expression level in *Ir8a^1^* flies. In *Ir25a^2^* flies, the IR genes that showed the greatest reduction in expression were those associated with humidity and temperature sensing (*Ir21a*, *Ir40a*, and *Ir93a*, but not *Ir68a*) (Figure 7A-B). In contrast, the only olfaction-associated tuning IR that was differentially expressed in *Ir25a^2^* or *Ir76b^1^* mutants was the amine-sensing receptor *Ir92a* in *Ir25a^2^* flies. Some of the other amine-sensing IRs had a trend towards reduced expression in *Ir25a^2^* or *Ir76b^1^* mutant lines, although this was not statistically significant: *Ir41a* (41% of control levels in *Ir25a^2^* flies) and *Ir75d* (75% and 63% of control levels in *Ir25a^2^* and *Ir76b^1^*, respectively). No IR receptors were differentially expressed in *Orco^2^* mutants.

**Figure 7:**
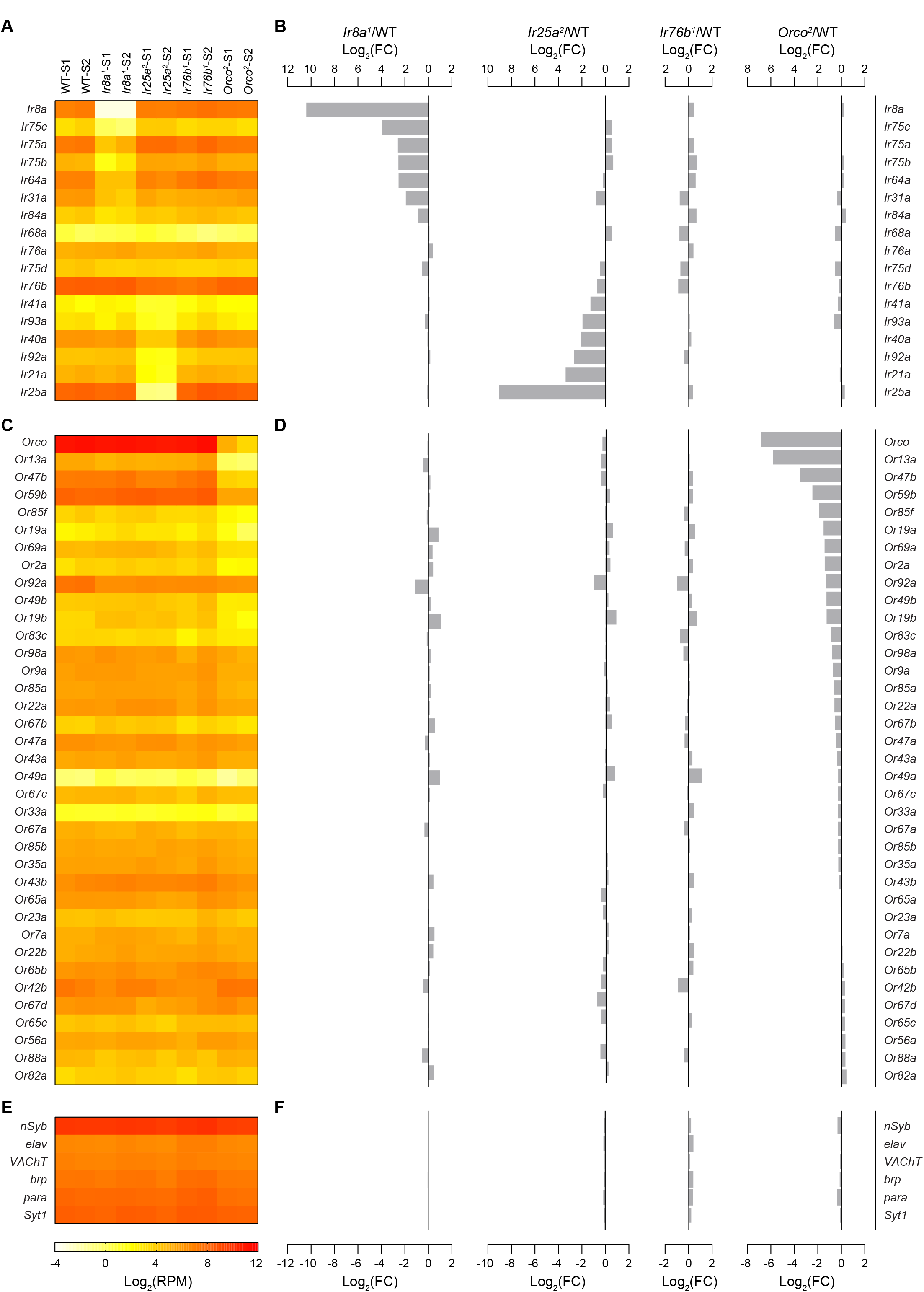
Expression of Ors/IRs changed selectively in co-receptor mutants. (A) A heat map showing the expression in Log_2_(RPM) of each of the known antennal IR genes in each of the ten samples. (B) Bar graphs reporting the log_2_ of the expression ratio between the co-receptor mutant and WT flies for *Ir8a^1^*, *Ir25a^2^*, *Ir76b^1^*, and *Orco^2^* flies. (C and D) Similar to (A) and (B), but for known antennal OR genes. (E and F) Similar to (A) and (B), but for a selection of neuronal markers.

Expression of many OR olfactory receptors was reduced in *Orco^2^* mutants (Figure 7C-D). Of the 36 reported antennal OR olfactory receptors, 10 were identified as *Orco^2^* DE genes, and an additional set of six OR receptors had a trend towards reduced expression with 35-68% of the expression level as controls (*Or19a*, *Or83c*, *Or98a*, *Or9a*, *Or85a*, and *Or22a*). There was no clear relationship among these 16 ORs with reduced expression, as they can be found in neurons in basiconic sensilla, trichoid and intermediate sensilla. In most sensilla subtypes, expression was reduced for the receptor in only one of its olfactory neurons, with equal likelihood of the OR associated with the large-spiking “A” neuron as with the small-spiking “B” neuron. This regulation of OR expression selectively occurred in *Orco^2^* flies, as there was only one OR gene that was differentially expressed in any of the three IR co-receptor mutants (*Or92a* in *Ir8a^1^* flies).

Finally, we wondered if the reduced olfactory receptor expression in the *Ir8a^1^*, *Ir25a^2^* and *Orco^2^* mutants might imply a broader loss of neurons in these flies. We examined the expression of six neuronal markers, and found that their expression was unaltered by the loss of co-receptors (Figure 7 E-F). Thus, there does not appear to be substantial neuronal degeneration in these flies at this age range (3-8 days post eclosion).

### Heat-responsive genes and *cyp450s* are enriched amongst *Ir25a^2^* DE genes

Unlike other olfactory receptor co-receptors, *Ir25a* has a unique additional role in humidity and temperature sensing [39–43]. This may explain why homozygous *Ir25a^2^* flies were somewhat sickly and more difficult to grow in large numbers compared to other olfactory co-receptor mutants.

These additional roles for *Ir25a* may also explain the large number of genes (90) that are differentially expressed in *Ir25a^2^* flies, but not in other co-receptor mutants (Figure 5C). We examined these genes in more detail, focusing on the 10 genes with the smallest p-value (Table 1). Excluding *Ir25a*, 3 of the 9 genes are reported to be involved in responding to heat: *Ir21a*, *GstE1*, and *Cyp6a17* [39, 44, 45]. Heat-associated genes among the other 80 DE genes include *jetboil* (*jtb*), a poorly characterized gene that contributes to thermal nociception, and *Ucp4C*, an oxidative phosphorylation uncoupling protein that mediates thermogenesis responses to cold temperatures [46, 47]. Further, seven of the nine genes were differentially expressed in *atonal* flies [48], which lack coeloconic olfactory sensilla, aristal thermosensory neurons, and hygrosensory and thermosensory sensilla in the sacculus [49].

**Table 1.** Top 10 genes only differentially expressed in *Ir25a^2^* mutants

| *Gene* | *Log<sub>2</sub>FC* | *Log<sub>2</sub>CPM* | *p-value* | *FDR* | *Description* |
| --- | --- | --- | --- | --- | --- |
| *Ir25a* | -9.32 | 7.79 | 5.74E-152 | 2.76E-148 | olfactory receptor |
| *CG10936* | -4.57 | 5.76 | 8.75E-75 | 1.68E-71 | no functional information |
| *Spn43Ab* | 3.54 | 5.67 | 9.51E-70 | 1.14E-66 | protease inhibitor |
| *Cyp4d21* | -3.12 | 9.95 | 3.99E-44 | 4.26E-41 | cytochrome p450, male enriched, mating |
| *ppk25* | -2.65 | 6.24 | 8.51E-40 | 7.43E-37 | DEG/ENaC ion channel, mating |
| *Ir21a* | -3.38 | 4.75 | 7.90E-35 | 6.33E-32 | ion channel, thermosensing |
| *Rgk2* | 4.18 | 3.56 | 9.85E-34 | 7.28E-31 | GTPase, calcium channel regulator |
| *GstE1* | 2.59 | 6.32 | 1.79E-32 | 1.15E-29 | glutathione transferase, response to heat |
| *phr* | 3.50 | 3.83 | 6.49E-31 | 3.54E-28 | DNA photolyase |
| *Cyp6a17* | -2.41 | 6.85 | 6.63E-31 | 3.54E-28 | cytochrome p450, temperature preference |

Members of the cytochrome p450 family were also enriched among the *Ir25a^2^* DE genes. Two, *Cyp4d21* and *Cyp6a17*, were among the top 10 DE genes in Table 1. Additionally, the GO term GO:0005506 “iron ion binding” was enriched amongst the genes downregulated in *Ir25a^2^* flies (Figure 6B), but not in other co-receptor mutants. These iron binding proteins are *Cyp4d21*, *Cyp6a17*, *Cyp9b1*, *Cyp6g2*, and *Cyp4p2* (Figure 8). All but *Cyp9b1* were also differentially expressed in *atonal* flies [48].

**Figure 8:**
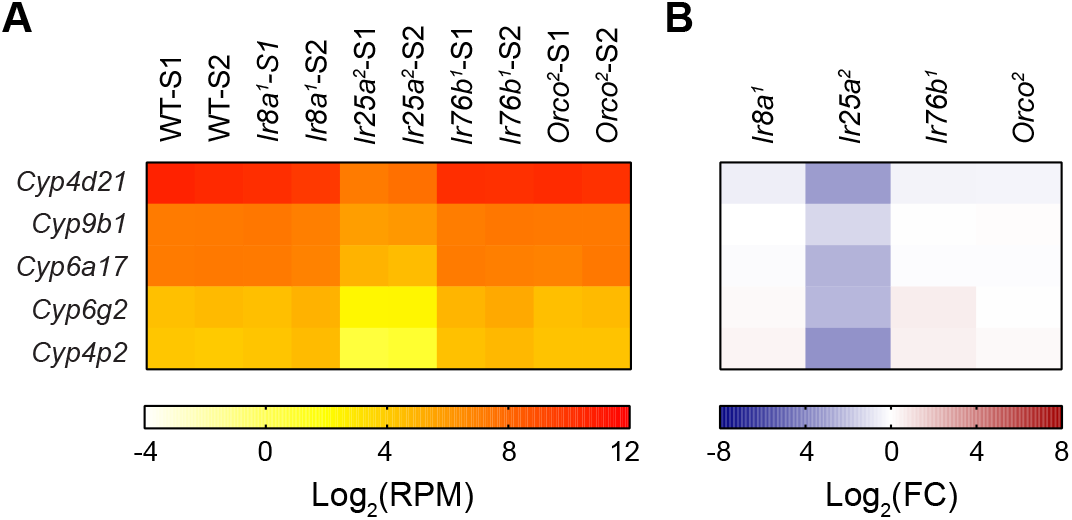
(A) A heat map showing the expression in Log_2_(RPM) in each of the ten samples of the Ir25a DE genes that are identified as “iron ion binding”. (B) A heat map reporting the log_2_ of the expression ratio between the co-receptor mutant and WT flies for *Ir8a^1^*, *Ir25a^2^*, *Ir76b^1^*, and *Orco^2^*.

## Discussion

This study clarifies and expands the roles of co-receptors in olfactory function in *Drosophila*. In particular, we demonstrate that both Ir76b and Ir25a function together as coreceptors for each of the tuning IR receptors that detect amines, and that the loss of Ir8a affects both the function and expression of acid-sensing tuning IRs.

Responses to amines are lost in ac2 and ac4 sensilla in mutants lacking either Ir25a or Ir76b expression (Figure 1). Phenethylamine responses in ac3B have a similar co-receptor dependence, implying that there is a currently unidentified tuning IR in ac3B ORNs and that each of the coeloconic sensilla has at least one acid-sensing and one amine-sensing neuron. We show that the co-receptor dependence of ac1 Ir92a ORNs is different in that the presence of either Ir25a or Ir76b fully maintains amine sensitivity, whereas responses are lost when both coreceptors are absent (Figure 2). This suggests that Ir92a can form functional receptors with either Ir25a or Ir76b. This is particularly intriguing because only Ir25a and Ir8a contain the aminoterminal domain involved in the assembly of IR subunits into heteromeric complexes [50, 51].

An overall picture is emerging of Ir25a and Ir76b often working together as co-receptors in chemosensory neurons. In addition to our findings in the olfactory system, studies using calcium imaging of gustatory neurons revealed that both co-receptors are necessary for responses to fatty acids, sour and carbonation [30–32]. However, other studies demonstrate that gustatory behaviors driven by amino acids, salt, and polyamines rely on Ir76b, but not Ir25a [28, 38, 52, 53]. Given that olfactory responses to polyamines show a similar selective dependence on Ir76b [28], despite additional Ir25a-dependence of responses in the polyamine sensitive Ir41a ORNs (Figure 1A), we believe that it would be informative to directly test the role of Ir25a in the responses of primary gustatory neurons to amino acids, salt, and polyamine.

Despite the similar roles of Ir76b and Ir25a as co-receptors for amine-sensing IRs, relatively few genes were differentially expressed in both genotypes (15 genes out of 199 *Ir25a^2^* DE genes and 87 *Ir76b^1^* DE genes, Figure 5C). Some of the *Ir25a^2^*-exclusive genes may contribute to its additional role in thermosensing, as indicated by several DE genes associated with heat sensitivity. These differential patterns of gene expression may also hint at unique roles for co-receptors beyond their roles in IR receptor function. For example, the expression of many *cyp450s* is reduced in *Ir25a* mutants, but not *Ir76b* mutants (Figure 8). Members of this family are best known for their roles in degradation of xenobiotics and metabolic biotransformation reactions, and they are thought to act as Odor Degrading Enzymes in the antenna. Additionally, eight transcription factors are among the 233 genes differentially expressed in at least one coreceptor mutant, and these may affect the expression of other genes in these flies. Most strikingly, expression of *twin-of-eyeless* (*toy*), a transcription factor known to play a role in nervous system development [54, 55], was reduced more than 100-fold in *Orco^2^* and *Ir25a^2^* flies and ~5 fold in *Ir76b^1^* flies (Dataset S1). Additionally, the neuronal transcription factor *castor* (*cas*) [56] was strongly upregulated in *Orco^2^* flies (Dataset S1). Given that many co-receptors are expressed by olfactory neurons in which they do not substantially contribute to odor responses, these DE genes may provide insight into additional functions of the co-receptors within neurons.

Our discovery that many OR receptors have reduced expression in *Orco^2^* flies is consistent with previous reports of progressive neuronal degeneration in the absence of Orco in *Drosophila*. The axons of some olfactory neurons degenerate over the first week post-eclosion in *Orco* mutants [15], and there is a ~30% reduction in the number of Or47 ORN cell bodies by the end of the second week [17]. Our data are the first to examine changes across all antennal ORs and reveal that the loss of OR expression varies dramatically, with >90% loss of *Or13a* and *Or47b* expression on one extreme and 15 receptors with <20% reduction on the other (Figure 7C-D). Similarly, maxillary palp Or46a ORNs are nearly eliminated in *Orco* mutants, whereas palp Or42a ORNs are unaffected [16], and loss of Orco in bees also differentially affects the expression of different OR tuning receptors [22]. Although there is evidence that this degeneration is due to the loss of neuronal activity in the absence of *Orco* [15], other factors must explain the large variability in susceptibility of different OR-expressing neurons, given that the odor-induced activity in each of these neurons depends on Orco. Further, the preservation of neuronal marker expression in Orco^2^ mutants suggests that reductions in olfactory receptor expression occur prior to or in the absence of neuronal cell death (Figure 7E-F).

We are the first to report a similar phenomenon in the absence of an IR co-receptor, finding greatly reduced expression of acid-sensing IRs in an *Ir8a* mutant (Figure 7A-B). Our data also reveal a similar effect on thermosensory and hygrosensory IR receptors in the absence of *Ir25a*. In contrast, expression of Ir25a- and Ir76b-dependent amine-sensing IRs was mostly unperturbed by the absence of either of these co-receptors, despite the associated loss of odor-sensitivity. One possibility is that Ir25a and Ir76b have a functionally redundant role in maintaining the expression of amine-sensing IRs.

Together our data establish Ir76b as an equal co-receptor to Ir25a in the olfactory system, and reveal the selective loss of OR and IR expression in certain co-receptor mutants. More broadly, the unique transcriptional changes induced by the loss of different co-receptors may provide clues to other roles for these genes in the function of the peripheral olfactory system.

## Materials and Methods

### Fly stocks

*Drosophila* flies were raised on a standard cornmeal-molasses-agar food in a 25°C incubator with a 12:12 hour light/dark cycle. Co-receptor mutant lines were obtained from the Bloomington Stock Center: #23130 *Orco^2^* [14], #41744 *Ir8a^1^* [12], #51309 *Ir76b^1^* [38], and #41737 *Ir25a^2^* [4]. Prior to experiments, each of these mutations was outcrossed for at least ten generations into *wCS*, our standard genetic background that is a *Cantonized w^1118^* line [57]. Such flies also served as the control flies (WT) for electrophysiology and RNASeq experiments.

### Electrophysiology

Single-sensillum recordings (SSR) were performed on 3-5 day old female flies as previously described [58, 59]. Flies were inserted into a trimmed 200 ul pipette tip, with a portion of the head and both antennae exposed. A tapered glass electrode stabilized one antenna against a glass coverslip. The prep was visualized under a BX51WI microscope (Olympus), and kept under a 2 L/min humidified air stream. A borosilicate reference electrode and an aluminosilicate recording electrode were filled with recording solution [60]. The reference electrode was inserted into an eye while the recording electrode was inserted into an individual sensillum with a MP-225 motorized micromanipulator (Sutter Instruments). An EXT-02F amplifier (NPI) with a custom 10x gain headstage was used for recording extracellular action potentials. A PowerLab 4/35 digitizer and LabChart Pro v8 software (ADInstruments) was used to acquire data at 10 kHz and AC filter at 300-1,700Hz.

Odorant cartridges were made pipetting 50 ul of odorant onto a 13 mm antibiotic assay disc (Whatman) inserted into a Pasteur pipette and closing the end of the pipette with a 1000 uL pipette tip. Odors were allowed to equilibrate for at least 20 minutes prior to experiments. Odors were applied by opening a Lee valve (02-21-08i) that allowed a 0.5 L/min air stream to travel through the cartridge and into the main air stream through a hole in the glass tube. Odorant delivery was controlled by a ValveBank 4 controller (Automate Scientific), which was driven by LabChart. Each LabChart trace was 10 s long, with a 1 s baseline, followed by 0.5 s odor application and 8.5s recovery period. An additional 10 s period was provided before application of another odorant to the same sensillum. Up to four sensilla were recorded per fly. Individual cartridges were allowed to recover for at least ten minutes between odorant applications and were used up to four times before being discarded.

Sensilla were recorded with a panel of ten odorants and two solvents. Propionic acid (99%, ACROS Organics, CAS 79-09-4), 1,4-diaminobutane (99%, ACROS Organics, CAS 110-60-1), pyrrolidine (>99%, ACROS Organics, CAS 123-75-1), trimethylamine (~45%, Sigma-Aldrich, CAS 75-50-3), and ammonium hydroxide (28-30%, Fisher, CAS 1336-21-6) were first diluted in water. Phenethylamine (99%, ACROS Organics, CAS 64-04-0), 1-hexanol (99%, ACROS Organics, CAS 111-27-3), 2-oxovaleric acid (>98%, Sigma-Aldrich, CAS 1821-02-9), phenylacetic acid (99%, Sigma-Aldrich, CAS 103-82-2), and phenylacetaldehyde (95%, Alfa Aesar, CAS 122-78-1) were diluted in paraffin oil. Odorants were first diluted to 10%, then diluted to 1% concentration, except for ammonium hydroxide and hexanol that were serially diluted to 0.1% and 0.001%, respectively.

Responses were analyzed using LabChart Spike Histogram module. Action potentials were counted over a 0.5 s response window, which was 100 ms after the stimulus onset due to the travel delay of the odorant in reaching the antenna. The action potentials of individual ORNs could not be sorted due to the similarity of their amplitudes, and therefore all spikes during the response period were counted. Odorant responses were calculated as responses to the odorant minus the responses to the solvent, either water or paraffin oil depending on the odorant. Sensillar classes were determined by their stereotypical location on the surface of the antenna and their responses to the odorant panel.

### RNA isolation and sequencing

Antennae were dissected from 3 to 8 day old flies, with approximately equal numbers of males and females. Two independent samples from wCS (WT), Ir25a^2^, Ir8a^1^, Ir76b^1^, and Orco^2^ flies were collected. Flies were dipped in liquid nitrogen, and antennae were manually dissected into 1.5 mL Eppendorf tubes in a liquid nitrogen bath. Antennae from 150-200 flies were dissected for each sample. A similar RNA isolation protocol was described previously [61]. Antennae were homogenized using disposable RNAse free micropestles and a QIAshredder column (Qiagen). RNA was extracted using an RNAeasy Micro Kit (Qiagen) and treated with DNAse from an iScript gDNA Clear cDNA Synthesis Kit (Bio-Rad). The Center for Genome Innovation at the University of Connecticut carried out quality control on the 10 RNA samples (~0.5 μg each) and generated libraries with the Illumina TruSeq Stranded mRNA library kit. Sequencing was conducted using an Illumina NextSeq 500 with 150 cycle output to produce ~24-40 million paired-end 75 bp reads per sample.

### RNASeq quantification and differential expression analysis

Raw reads were stored in BaseSpace Sequence Hub (Illumina) and downloaded to the University of Connecticut High Performance Computing cluster. Trimmed reads were generated using Sickle with the default parameters [62]. The *Drosophila* gene annotation file and reference genome were downloaded from Ensembl (BDGP6) [63]. Trimmed reads were aligned to the reference genome with mitochondrial genes removed using STAR with the default parameters [64]. Mapped reads generated by STAR are available under BioProject accession number PRJNA735732 at the NCBI Short Read Archive (SRA). Raw counts of reads mapping uniquely to each gene were generated using HTSeq [65]. HTSeq-count was used with “intersection-nonempty” and “nonunique-none” modes to handle reads mapping to more than one feature. To analyze expression of Ir76b on an exon-by-exon basis, we fed the aligned reads from STAR into DEXSeq and ran the program with the default parameters to estimate the RPM for each exon of each gene [66, 67].

Differential gene expression analysis was carried out on the raw gene counts from HTSeq-count with the EdgeR v3.8 package in R studio v3.6.1 (Robinson et al., 2010). First, EdgeR was used to generate a general linearized model (GLM) over all ten samples and identified 416 genes that were differentially expressed with a FDR < 0.05. To then identify the specific samples in which the genes were differentially expressed with a FDR < 0.05, the classic EdgeR (ExactTest) was run between samples for each co-receptor and the WT samples in a pair-wise fashion. We identified DE genes for each mutant as those with FDR < 0.05 and with a >2-fold change in expression. The 233 DE genes that were those that were identified as DE with the ExactTest and GLM in at least one genotype. The Reads per Million Mapped Reads (RPM) value for each gene in each sample was calculated by EdgeR. To calculate the Log_2_(RPM) values and Log_2_(FC) in graphs in Figures 4–8, Tables 1-4, and Table S1, a small value was added to each RPM value to avoid undefined fold-changes when one sample has zero reads.

The identification of significantly enriched Gene Ontology (GO) terms was carried out with PANTHER Overrepresentation Tests run through the AmiGO2 website using the GO Ontology Database released 01-01-2021 [68–71]. Separate analyses were run for upregulated and downregulated DE genes for each co-receptor mutant. The Fisher’s Exact test was run with the False Discovery Rate correction for multiple comparisons.

### Statistical Analysis

Data were analyzed using GraphPad Prism 9. Box plots depict the median and interquartile range, and the whiskers span the minimum to maximum data points. All max individual data points are overlaid on the box plots. Mann-Whitney tests were used to compare two genotypes. For comparisons of four genotypes, a Kruskal-Wallis test was used and the FDR was controlled with the two-stage linear step up method of Benjamini, Krieger, and Yekutieli. Data Statistical parameters can be found in the figure legends. Genotypes that are significantly different (p < 0.05) are indicated with different letters above the box plots.

## Acknowledgements

We acknowledge the Bloomington *Drosophila* Stock Center (NIH P40OD018537) for fly lines. We thank Eleanor LoConte for assistance with outcrossing the mutants and Ben Groce for help with antennal dissections. We thank Chih-Ying Suh, Chris Potter, and Anastasios Tzingounis for discussions and comments on the manuscript. This research was supported by NIH awards R21DC017868 and R35GM133209 to K.M.

## Dataset S1

List of all differentially expressed genes in each genotype. This dataset contains five spreadsheets. The first four list the differentially expressed genes in each genotype, with analysis for each co-receptor mutants on a separate spreadsheet. On the left of each spreadsheet are the data from the EdgeR ExactTest run against the WT control in yellow. The first column contains the gene symbol, and is followed by a column with the log of fold-change in gene expression (log_2_(RPM_mutant_/RPM_WT_). The third column contains each gene’s average expression across the four samples (log_2_(RPM)). The following two columns list the p-value and false discovery rate (FDR) for differential expression of the gene between the two genotypes. On the right in orange is the expression of each gene in each sample in RPM. The fifth spreadsheet “233 DE Genes” is a summary of the DE genes across genotypes. The DE genes are listed in the first column by their gene symbol. The next four columns identifies whether the gene is differentially expressed in each of the four co-receptor mutants, with differentially expressed genes marked by an X. Columns G through P report the expression level (RPM) of each gene in each of the two samples from each of the five genotypes.

## Notes

### Competing Interest Statement

The authors have declared no competing interest.

## References

1. Rytz, R., Croset, V., and Benton, R. (2013). Ionotropic receptors (IRs): chemosensory ionotropic glutamate receptors in *Drosophila* and beyond. Insect Biochem Mol Biol 43, 888–897.

2. Vosshall, L.B., and Stocker, R.F. (2007). Molecular architecture of smell and taste in *Drosophila*. Annu. Rev. Neurosci. 30, 505–533.

3. Silbering, A.F., Rytz, R., Grosjean, Y., Abuin, L., Ramdya, P., Jefferis, G.S., and Benton, R. (2011). Complementary function and integrated wiring of the evolutionarily distinct *Drosophila* olfactory subsystems. J. Neurosci. 31, 13357–13375.

4. Benton, R., Vannice, K.S., Gomez-Diaz, C., and Vosshall, L.B. (2009). Variant Ionotropic Glutamate Receptors as Chemosensory Receptors in *Drosophila*. Cell 136, 149–162.

5. Couto, A., Alenius, M., and Dickson, B.J. (2005). Molecular, anatomical, and functional organization of the Drosophila olfactory system. Curr. Biol. 15, 1535–1547.

6. Fishilevich, E., and Vosshall, L.B. (2005). Genetic and functional subdivision of the Drosophila antennal lobe. Curr. Biol. 15, 1548–1553.

7. Hallem, E.A., Dahanukar, A., and Carlson, J.R. (2006). Insect odor and taste receptors. Annu. Rev. Entomol. 51, 113–135.

8. Kurtovic, A., Widmer, A., and Dickson, B.J. (2007). A single class of olfactory neurons mediates behavioural responses to a Drosophila sex pheromone. Nature 446, 542–546.

9. Dweck, H.K., Ebrahim, S.A., Thoma, M., Mohamed, A.A., Keesey, I.W., Trona, F., Lavista-Llanos, S., Svatoš, A., Sachse, S., and Knaden, M. (2015). Pheromones mediating copulation and attraction in Drosophila. Proc. Natl. Acad. Sci. 112, E2829–E2835.

10. Lin, H.H., Cao, D.S., Sethi, S., Zeng, Z., Chin, J.S.R., Chakraborty, T.S., Shepherd, A.K., Nguyen, C.A., Yew, J.Y., Su, C.Y., et al. (2016). Hormonal Modulation of Pheromone Detection Enhances Male Courtship Success. Neuron 90, 1272–1285.

11. Sato, K., Pellegrino, M., Nakagawa, T., Nakagawa, T., Vosshall, L.B., and Touhara, K. (2008). Insect olfactory receptors are heteromeric ligand-gated ion channels. Nature 452, 1002–1006.

12. Abuin, L., Bargeton, B., Ulbrich, M.H., Isacoff, E.Y., Kellenberger, S., and Benton, R. (2011). Functional architecture of olfactory ionotropic glutamate receptors. Neuron 69, 44–60.

13. Butterwick, J.A., Del Marmol, J., Kim, K.H., Kahlson, M.A., Rogow, J.A., Walz, T., and Ruta, V. (2018). Cryo-EM structure of the insect olfactory receptor Orco. Nature 560, 447–452.

14. Larsson, M.C., Domingos, A.I., Jones, W.D., Chiappe, M.E., Amrein, H., and Vosshall, L.B. (2004). Or83b encodes a broadly expressed odorant receptor essential for *Drosophila* olfaction. Neuron 43, 703–714.

15. Chiang, A., Priya, R., Ramaswami, M., Vijayraghavan, K., and Rodrigues, V. (2009). Neuronal activity and Wnt signaling act through Gsk3-beta to regulate axonal integrity in mature Drosophila olfactory sensory neurons. Development 136, 1273–1282.

16. Task, D., and Potter, C.J. (2021). Rapid degeneration of Drosophila olfactory neurons in Orco mutant maxillary palps. microPublication Biology 2021.

17. Hueston, C.E., Olsen, D., Li, Q., Okuwa, S., Peng, B., Wu, J., and Volkan, P.C. (2016). Chromatin Modulatory Proteins and Olfactory Receptor Signaling in the Refinement and Maintenance of Fruitless Expression in Olfactory Receptor Neurons. PLoS Biol. 14, e1002443.

18. DeGennaro, M., McBride, C.S., Seeholzer, L., Nakagawa, T., Dennis, E.J., Goldman, C., Jasinskiene, N., James, A.A., and Vosshall, L.B. (2013). orco mutant mosquitoes lose strong preference for humans and are not repelled by volatile DEET. Nature 498, 487–491.

19. Li, Y., Zhang, J., Chen, D., Yang, P., Jiang, F., Wang, X., and Kang, L. (2016). CRISPR/Cas9 in locusts: Successful establishment of an olfactory deficiency line by targeting the mutagenesis of an odorant receptor co-receptor (Orco). Insect Biochem Mol Biol 79, 27–35.

20. Liu, Q., Liu, W., Zeng, B., Wang, G., Hao, D., and Huang, Y. (2017). Deletion of the Bombyx mori odorant receptor co-receptor (BmOrco) impairs olfactory sensitivity in silkworms. Insect Biochem Mol Biol 86, 58–67.

21. Yan, H., Opachaloemphan, C., Mancini, G., Yang, H., Gallitto, M., Mlejnek, J., Leibholz, A., Haight, K., Ghaninia, M., and Huo, L. (2017). An engineered orco mutation produces aberrant social behavior and defective neural development in ants. Cell 170, 736–747. e739.

22. Chen, Z., Traniello, I.M., Rana, S., Cash-Ahmed, A.C., Sankey, A.L., Yang, C., and Robinson, G.E. (2021). Neurodevelopmental and transcriptomic effects of CRISPR/Cas9-induced somatic orco mutation in honey bees. J. Neurogenet., 1–13.

23. Trible, W., Olivos-Cisneros, L., McKenzie, S.K., Saragosti, J., Chang, N.-C., Matthews, B.J., Oxley, P.R., and Kronauer, D.J. (2017). Orco mutagenesis causes loss of antennal lobe glomeruli and impaired social behavior in ants. Cell 170, 727–735. e710.

24. Yao, C.A., Ignell, R., and Carlson, J.R. (2005). Chemosensory coding by neurons in the coeloconic sensilla of the *Drosophila* antenna. J. Neurosci. 25, 8359–8367.

25. Ai, M., Blais, S., Park, J.Y., Min, S., Neubert, T.A., and Suh, G.S. (2013). Ionotropic glutamate receptors IR64a and IR8a form a functional odorant receptor complex in vivo in Drosophila. J. Neurosci. 33, 10741–10749.

26. Raji, J.I., Melo, N., Castillo, J.S., Gonzalez, S., Saldana, V., Stensmyr, M.C., and DeGennaro, M. (2019). Aedes aegypti Mosquitoes Detect Acidic Volatiles Found in Human Odor Using the IR8a Pathway. Curr. Biol. 29, 1253–1262 e1257.

27. Zhang, J., Bisch-Knaden, S., Fandino, R.A., Yan, S., Obiero, G.F., Grosse-Wilde, E., Hansson, B.S., and Knaden, M. (2019). The olfactory coreceptor IR8a governs larval feces-mediated competition avoidance in a hawkmoth. Proc Natl Acad Sci U S A 116, 21828–21833.

28. Hussain, A., Zhang, M., Ucpunar, H.K., Svensson, T., Quillery, E., Gompel, N., Ignell, R., and Grunwald Kadow, I.C. (2016). Ionotropic chemosensory receptors mediate the taste and smell of polyamines. PLoS Biol. 14, e1002454.

29. Min, S., Ai, M., Shin, S.A., and Suh, G.S. (2013). Dedicated olfactory neurons mediating attraction behavior to ammonia and amines in *Drosophila*. Proc. Natl. Acad. Sci. USA 110, E1321–1329.

30. Chen, Y., and Amrein, H. (2017). Ionotropic receptors mediate Drosophila oviposition preference through sour gustatory receptor neurons. Curr. Biol. 27, 2741–2750. e2744.

31. Ahn, J.-E., Chen, Y., and Amrein, H. (2017). Molecular basis of fatty acid taste in Drosophila. Elife 6, e30115.

32. Sanchez-Alcaniz, J.A., Silbering, A.F., Croset, V., Zappia, G., Sivasubramaniam, A.K., Abuin, L., Sahai, S.Y., Munch, D., Steck, K., Auer, T.O., et al. (2018). An expression atlas of variant ionotropic glutamate receptors identifies a molecular basis of carbonation sensing. Nat. Commun. 9, 4252.

33. Younger, M.A., Herre, M., Ehrlich, A.R., Gong, Z., Gilbert, Z.N., Rahiel, S., Matthews, B.J., and Vosshall, L.B. (2020). Non-canonical odor coding ensures unbreakable mosquito attraction to humans. bioRxiv, 10.1101/2020.1111.1107.368720v368721.

34. Task, D., Lin, C.-C., Afify, A., Li, H., Vulpe, A., Menuz, K., and Potter, C.J. (2020). Widespread Polymodal Chemosensory Receptor Expression in Drosophila Olfactory Neurons. bioRxiv, 10.1101/2020.1111.1107.355651.

35. Vulpe, A., Kim, H.S., Ballou, S., Wu, S.T., Grabe, V., Nava Gonzales, C., Liang, T., Sachse, S., Jeanne, J.M., Su, C.Y., et al. (2021). An ammonium transporter is a non-canonical olfactory receptor for ammonia. Curr. Biol., Forthcoming.

36. Prieto-Godino, L.L., Rytz, R., Cruchet, S., Bargeton, B., Abuin, L., Silbering, A.F., Ruta, V., Dal Peraro, M., and Benton, R. (2017). Evolution of Acid-Sensing Olfactory Circuits in Drosophilids. Neuron 93, 661–676 e666.

37. Grosjean, Y., Rytz, R., Farine, J.P., Abuin, L., Cortot, J., Jefferis, G.S., and Benton, R. (2011). An olfactory receptor for food-derived odours promotes male courtship in Drosophila. Nature 478, 236–240.

38. Zhang, Y.V., Ni, J., and Montell, C. (2013). The molecular basis for attractive salt-taste coding in Drosophila. Science 340, 1334–1338.

39. Budelli, G., Ni, L., Berciu, C., van Giesen, L., Knecht, Z.A., Chang, E.C., Kaminski, B., Silbering, A.F., Samuel, A., Klein, M., et al. (2019). Ionotropic Receptors Specify the Morphogenesis of Phasic Sensors Controlling Rapid Thermal Preference in Drosophila. Neuron 101, 738–747.e733.

40. Knecht, Z.A., Silbering, A.F., Ni, L., Klein, M., Budelli, G., Bell, R., Abuin, L., Ferrer, A.J., Samuel, A.D., and Benton, R. (2016). Distinct combinations of variant ionotropic glutamate receptors mediate thermosensation and hygrosensation in Drosophila. Elife 5, e17879.

41. Ni, L., Klein, M., Svec, K.V., Budelli, G., Chang, E.C., Ferrer, A.J., Benton, R., Samuel, A.D., and Garrity, P.A. (2016). The ionotropic receptors IR21a and IR25a mediate cool sensing in Drosophila. Elife 5, e13254.

42. Enjin, A., Zaharieva, E.E., Frank, D.D., Mansourian, S., Suh, G.S., Gallio, M., and Stensmyr, M.C. (2016). Humidity sensing in Drosophila. Curr. Biol. 26, 1352–1358.

43. Knecht, Z.A., Silbering, A.F., Cruz, J., Yang, L., Croset, V., Benton, R., and Garrity, P.A. (2017). Ionotropic Receptor-dependent moist and dry cells control hygrosensation in Drosophila. Elife 6, e26654.

44. Neal, S.J., Karunanithi, S., Best, A., So, A.K., Tanguay, R.M., Atwood, H.L., and Westwood, J.T. (2006). Thermoprotection of synaptic transmission in a Drosophila heat shock factor mutant is accompanied by increased expression of Hsp83 and DnaJ-1. Physiol. Genomics 25, 493–501.

45. Kang, J., Kim, J., and Choi, K.W. (2011). Novel cytochrome P450, cyp6a17, is required for temperature preference behavior in Drosophila. PLoS One 6, e29800.

46. Honjo, K., Mauthner, S.E., Wang, Y., Skene, J.H.P., and Tracey, W.D., Jr. (2016). Nociceptor-Enriched Genes Required for Normal Thermal Nociception. Cell Rep. 16, 295–303.

47. Da-Re, C., De Pitta, C., Zordan, M.A., Teza, G., Nestola, F., Zeviani, M., Costa, R., and Bernardi, P. (2014). UCP4C mediates uncoupled respiration in larvae of Drosophila melanogaster. EMBO Rep 15, 586–591.

48. Menuz, K., Larter, N.K., Park, J., and Carlson, J.R. (2014). An RNA-seq screen of the *Drosophila* antenna identifies a transporter necessary for ammonia detection. PLoS Genet. 10, e1004810.

49. Gupta, B.P., and Rodrigues, V. (1997). Atonal is a proneural gene for a subset of olfactory sense organs in *Drosophila*. Genes Cells 2, 225–233.

50. Croset, V., Rytz, R., Cummins, S.F., Budd, A., Brawand, D., Kaessmann, H., Gibson, T.J., and Benton, R. (2010). Ancient protostome origin of chemosensory ionotropic glutamate receptors and the evolution of insect taste and olfaction. PLoS Genet. 6, e1001064.

51. Abuin, L., Prieto-Godino, L.L., Pan, H., Gutierrez, C., Huang, L., Jin, R., and Benton, R. (2019). In vivo assembly and trafficking of olfactory Ionotropic Receptors. BMC Biol. 17, 34.

52. Ganguly, A., Pang, L., Duong, V.-K., Lee, A., Schoniger, H., Varady, E., and Dahanukar, A. (2017). A molecular and cellular context-dependent role for Ir76b in detection of amino acid taste. Cell Rep. 18, 737–750.

53. Croset, V., Schleyer, M., Arguello, J.R., Gerber, B., and Benton, R. (2016). A molecular and neuronal basis for amino acid sensing in the Drosophila larva. Sci. Rep. 6, 34871.

54. Kurusu, M., Nagao, T., Walldorf, U., Flister, S., Gehring, W.J., and Furukubo-Tokunaga, K. (2000). Genetic control of development of the mushroom bodies, the associative learning centers in the Drosophila brain, by the eyeless, twin of eyeless, and Dachshund genes. Proc Natl Acad Sci U S A 97, 2140–2144.

55. Czerny, T., Halder, G., Kloter, U., Souabni, A., Gehring, W.J., and Busslinger, M. (1999). twin of eyeless, a second Pax-6 gene of Drosophila, acts upstream of eyeless in the control of eye development. Mol. Cell 3, 297–307.

56. Hitier, R., Chaminade, M., and Preat, T. (2001). The Drosophila castor gene is involved in postembryonic brain development. Mech. Dev. 103, 3–11.

57. Koh, T.W., He, Z., Gorur-Shandilya, S., Menuz, K., Larter, N.K., Stewart, S., and Carlson, J.R. (2014). The Drosophila IR20a clade of ionotropic receptors are candidate taste and pheromone receptors. Neuron 83, 850–865.

58. Benton, R., and Dahanukar, A. (2011). Electrophysiological recording from Drosophila taste sensilla. Cold Spring Harb Protoc 2011, 839–850.

59. Dobritsa, A.A., van der Goes van Naters, W., Warr, C.G., Steinbrecht, R.A., and Carlson, J.R. (2003). Integrating the molecular and cellular basis of odor coding in the Drosophila antenna. Neuron 37, 827–841.

60. Kaissling, K.E., and Thorson, J. (1980). Insect olfactory sensilla: Structural, chemical and electrical aspects of the functional organization. In Receptors for neurotransmitters, hormones, and pheromones in insects, D.B. Sattelle, L.M. Hall and J.G. Hildebrand, eds. (Amsterdam: Elsevier), pp. 261–282.

61. Mohapatra, P., and Menuz, K. (2019). Molecular Profiling of the Drosophila Antenna Reveals Conserved Genes Underlying Olfaction in Insects. G3 (Bethesda) 9, 3753–3771.

62. Joshi, N.A., and Fass, J.N. (2011). Sickle: A sliding-window, adaptive, quality-based trimming tool for FastQ files. 1.33 Edition.

63. Yates, A.D., Achuthan, P., Akanni, W., Allen, J., Allen, J., Alvarez-Jarreta, J., Amode, M.R., Armean, I.M., Azov, A.G., Bennett, R., et al. (2020). Ensembl 2020. Nucleic Acids Res. 48, D682–D688.

64. Dobin, A., Davis, C.A., Schlesinger, F., Drenkow, J., Zaleski, C., Jha, S., Batut, P., Chaisson, M., and Gingeras, T.R. (2013). STAR: ultrafast universal RNA-seq aligner. Bioinformatics 29, 15–21.

65. Anders, S., Pyl, P.T., and Huber, W. (2015). HTSeq—a Python framework to work with high-throughput sequencing data. Bioinformatics 31, 166–169.

66. Anders, S., Reyes, A., and Huber, W. (2012). Detecting differential usage of exons from RNA-seq data. Genome Res. 22, 2008–2017.

67. Reyes, A., Anders, S., Weatheritt, R.J., Gibson, T.J., Steinmetz, L.M., and Huber, W. (2013). Drift and conservation of differential exon usage across tissues in primate species. Proc Natl Acad Sci U S A 110, 15377–15382.

68. Mi, H., Muruganujan, A., Ebert, D., Huang, X., and Thomas, P.D. (2019). PANTHER version 14: more genomes, a new PANTHER GO-slim and improvements in enrichment analysis tools. Nucleic Acids Res. 47, D419–D426.

69. Carbon, S., Ireland, A., Mungall, C.J., Shu, S., Marshall, B., and Lewis, S. (2009). AmiGO: online access to ontology and annotation data. Bioinformatics 25, 288–289.

70. Gene Ontology Consortium (2021). The Gene Ontology resource: enriching a GOld mine. Nucleic Acids Res. 49, D325–D334.

71. Ashburner, M., Ball, C.A., Blake, J.A., Botstein, D., Butler, H., Cherry, J.M., Davis, A.P., Dolinski, K., Dwight, S.S., and Eppig, J.T. (2000). Gene ontology: tool for the unification of biology. Nat. Genet. 25, 25.

